# APOBEC3 antagonism fully explains HIV-1 Vif essentiality under interferon and differentiation conditions

**DOI:** 10.64898/2026.04.16.719124

**Authors:** Ryo Shimizu, Michael Jonathan, Hiromi Terasawa, Akatsuki Saito, Kazuaki Monde, Terumasa Ikeda

## Abstract

HIV-1 virion infectivity factor (Vif) counteracts APOBEC3 (A3) proteins by targeting them for proteasomal degradation, thereby preventing lethal G-to-A mutations and loss of infectivity. However, Vif also degrades additional cellular proteins, raising the possibility that its essential role in infectious virion production may extend beyond A3 antagonism, particularly under inflammatory or differentiation conditions. Whether such conditions reveal additional essential Vif targets remains unresolved. Here, using interferon (IFN)-stimulated or phorbol 12-myristate 13-acetate (PMA)-differentiated THP-1 monocytic cells, we directly addressed this question. Although type I IFN and PMA markedly suppressed viral production through Vif-independent mechanisms, ΔVif viruses produced from stimulated parental cells exhibited severely reduced infectivity. In contrast, disruption of A3A-A3G fully restored ΔVif infectivity to wild-type levels under all conditions tested. G-to-A mutations were attributable exclusively to A3 proteins under both IFN and PMA stimulation, whereas IFN-induced, A3-independent blocks to reverse transcription were not antagonized by Vif. Across diverse HIV-1 strains, the requirement for Vif was strictly dependent on A3 family proteins. These findings demonstrate that the essential role of Vif is fully explained by antagonism of A3-mediated restriction and that evolutionary pressure on Vif is driven predominantly by the need to counteract A3-mediated restriction and mutagenesis.

**Importance:** A3 proteins are potent intrinsic antiviral factors that restrict HIV-1 by inducing lethal mutations and impairing reverse transcription. The viral protein Vif counteracts A3-mediated restriction, yet it also targets additional host proteins, raising questions about whether its essential role extends beyond A3 antagonism under inflammatory conditions. Myeloid cells mount IFN responses and undergo transcriptional reprogramming upon differentiation, environments in which additional Vif functions could emerge. Using an A3A-A3G-null monocytic cell model, we show that even under IFN stimulation or PMA-induced differentiation, the requirement for Vif in maintaining HIV-1 infectivity is due to A3 proteins. In contrast, Vif does not counteract IFN-induced, A3-independent restriction pathways, highlighting functional specificity. These findings establish the A3-Vif axis as the central determinant of Vif dependency and indicate that evolutionary pressure on Vif is driven predominantly by the need to evade A3-mediated restriction. Targeting Vif may therefore expose HIV-1 to intrinsic inactivation, even in inflammatory environments.

## Introduction

The Apolipoprotein B mRNA editing enzyme catalytic polypeptide-like 3 (APOBEC3, A3) family of cytosine deaminases catalyzes C-to-U deamination in single-stranded DNA and functions as a potent intrinsic restriction system against retroviruses, particularly HIV-1 [reviewed in (1–3)]. In humans, seven *A3* genes (*A3A*-*A3D* and *A3F*-*A3H*) are clustered on chromosome 22 between *CBX6* and *CBX7* genes and encode proteins containing either single (A3A, A3C, and A3H) or double (A3B, A3D, A3F, and A3G) zinc-binding domains [reviewed in (1–6)]. Up to five A3 proteins (A3C-I188, A3D, A3F, A3G, and stable A3H haplotypes) contribute to HIV-1 restriction in CD4+ T cells and myeloid cells (7–17).

A3 proteins employ both deaminase-dependent and -independent mechanisms [reviewed in (1–3, 18, 19)]. A3G preferentially deaminates cytosines within the 5’-C<u>C</u>-3’ motifs (the underlined C is the target C), whereas other A3 proteins favor 5’-T<u>C</u>-3’ contexts (7, 8, 10, 15, 16, 20–26). Deaminase-independent mechanisms include the roadblock model, in which A3 proteins impede reverse transcription, as well as direct interactions with the Reverse Transcriptase (16, 27–30). These non-catalytic mechanisms appear to be particularly important for A3F and A3H (31–34). Through these complementary activities, A3 enzymes impose potent mutational and replicative barriers on HIV-1.

To counteract this antiviral system, HIV-1 encodes the virion infectivity factor (Vif), which recruits a host E3 ubiquitin ligase complex (CBF-β, CUL5, ELOB, ELOC, RBX2, and ARIH2) to target A3 proteins for proteasomal degradation (35–44). A3-mediated mutagenesis imposes strong selective pressure on lentiviral genomes, underscoring the central evolutionary role of Vif. In addition to degrading A3 proteins, Vif has been reported to target other cellular factors, including PPP2R5 family members, DPH7 and FMRP (35, 45–50). Although the contribution of these proteins to infectious virion production remains incompletely defined, these observations raise a fundamental question: does Vif possess essential cellular targets beyond A3 proteins?

This question is particularly relevant in myeloid cells, which exhibit robust type I interferon (IFN) responses and express high levels of interferon-stimulated antiviral factors, including A3 family members as well as BST2/Tetherin, TRIM5α, IFITMs, MX2, and GBP5 (51–60). IFN stimulation induces a broad antiviral transcriptional program (61–63). In addition, phorbol 12-myristate 13-acetate (PMA) treatment drives macrophage-like differentiation and extensive transcriptional remodeling in THP-1 cells (53, 56, 62–65). These inflammatory and differentiation signals could potentially expose context-dependent Vif functions not apparent under basal conditions.

We previously demonstrated that A3 proteins are the primary Vif targets required for infectious HIV-1 production in THP-1 monocytic cells (16). However, it has remained unclear whether Vif possesses additional essential functions under inflammatory or differentiation conditions. Here, we directly address this question by using A3A-A3G-null THP-1 cells to determine whether Vif has any indispensable targets other than A3 proteins under IFN-stimulated or PMA-differentiated conditions.

## Results

### THP-1 parental and A3A-A3G-null THP-1 cells exhibit comparable responsiveness to type I IFN and PMA

Several A3 family members are upregulated with type I IFN treatment (16, 53–55, 65–68). In addition, PMA induces differentiation of THP-1 cells from non-adherent suspension cells into macrophage-like adherent cells and alters both the transcriptional landscape (62, 63) and antiviral phenotype (53, 56, 64). To validate Vif function under stimulated conditions, we confirmed that parental THP-1 and A3A-A3G-null cells (clone #11-4) responded comparably to IFN and PMA treatment by reverse transcription quantitative polymerase chain reaction (RT-qPCR).

As expected, IFN robustly induced *A3A* mRNA (∼100-fold) and moderately upregulated *A3F, A3G,* and *A3H* mRNAs (3- to 4-fold) in parental THP-1 cells, whereas *A3B, A3C,* and *A3D* were only slightly induced (< 1.5-fold) (**Fig. 1A**). Consistent with previous findings (16), *A3B, A3C, A3D, A3F,* and *A3G* mRNAs remained undetectable in THP-1#11-4 cells, while *A3A* and *A3H* mRNAs were retained (**Fig. 1A**). Notably, A3A protein is not expressed in this clone even after IFN treatment, and both parental and THP-1#11-4 cells express A3H haplotype I, which encodes an unstable and functionally inactive protein with limited antiviral activity against HIV-1 (16).

**Figure 1.**
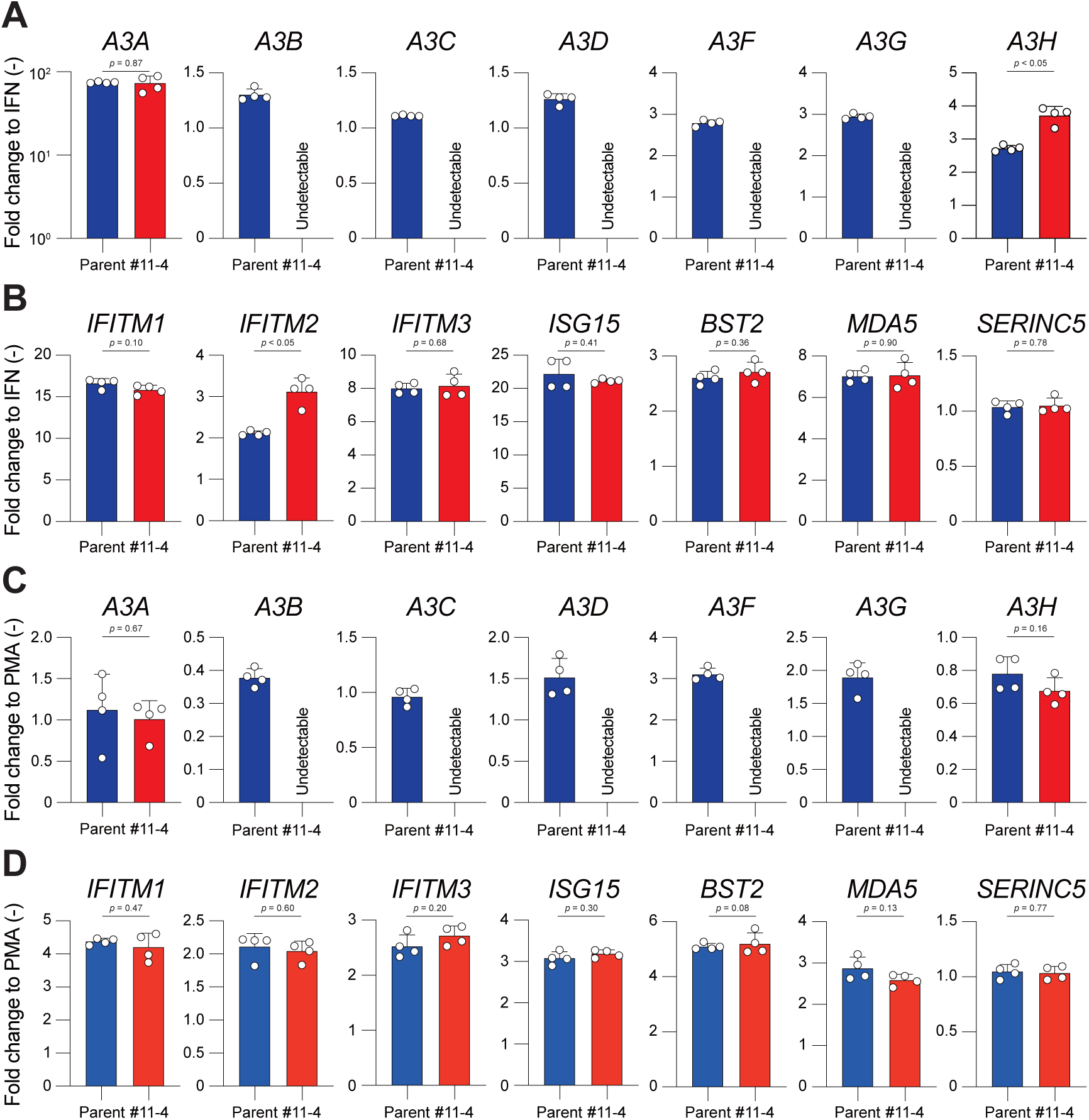
mRNA expression levels of *A3* family members and ISGs in response to type I IFN or PMA treatment. **(A)** RT-qPCR analysis of *A3* mRNAs following type I IFN treatment. Parental and THP-1 #11-4 cells were treated with 500 units/ml type I IFN for 6 hours. Total RNA was extracted and *A3* mRNA levels were quantified by RT-qPCR, normalized to *TATA-binding protein* (*TBP*) mRNA, and expressed as fold change relative to the untreated cells. Bars represent mean ± standard deviation (SD) of four technical replicates from one representative experiment (n = 3 independent experiments). Statistical significance between parental and THP-1#11-4 cell was assessed using an unpaired Welch’s t-test; p-values are indicated. **(B)** RT-qPCR analysis of the indicated ISGs following type I IFN treatment. Cells were treated and analyzed as in (**A**). Bars represent mean ± SD of four technical replicates from one representative experiment (n = 3 independent experiments). Statistical analysis was performed as described above. **(C)** RT-qPCR analysis of *A3* mRNAs following PMA treatment. Parental and THP-1 #11-4 cells were treated with 20 ng/ml PMA for 20 hours. RNA extraction and analysis were performed as in (**A**). Bars represent mean ± SD of four technical replicates from one representative experiment (n = 3 independent experiments). Statistical significance was assessed as described above. **(D)** RT-qPCR analysis of the indicated ISGs following PMA treatment. Cells were treated and analyzed as in (**C**). Bars represent mean ± SD of three independent experiments. Statistical analysis was performed as described above.

Because IFN induces numerous interferon-stimulated genes (ISGs) beyond the A3 family proteins [reviewed in (69–71)], we next assessed broader IFN responsiveness. IFN treatment induced *IFITM1*, *IFITM2*, *IFITM3*, *ISG15*, *BST2*, *MDA5*, and *SERINC5* mRNAs to comparable levels in both cell lines, with only minor differences (slightly higher *IFITM2* induction in THP-1#11-4 cells and no induction of *SERINC5* in either line) (**Fig. 1B**).

Under PMA treatment, *A3F* and *A3G* mRNAs were moderately increased (2- to 3-fold) in parental THP-1 cells, whereas other *A3* genes showed only modest changes (0.8- to 1.5-fold), with *A3B* mRNA reduced to ∼0.4-fold (**Fig. 1C**). As with IFN treatment, *A3B, A3C, A3D, A3F,* and *A3G* transcripts remained undetectable in THP-1#11-4 cells following PMA treatment, while *A3A* and *A3H* mRNA levels were comparable between the two cell lines (**Fig. 1A**, **1C**). PMA also induced representative ISGs (2- to 5-fold) in both cell types, although to a lesser extent than IFN, and *SERINC5* remained unchanged (**Fig. 1D**).

Collectively, these data demonstrate that parental and THP-1#11-4 cells exhibit comparable transcriptional responsiveness to both IFN and PMA stimulation, validating THP-1#11-4 cells as an appropriate system to dissect Vif-dependent effects under stimulated and differentiated conditions.

### IFN and PMA suppress HIV-1 production independent of Vif and A3 proteins

We next examined whether IFN or PMA stimulation alters Vif-dependent viral production. Pseudo-single cycle infectivity assays were performed as previously described (15, 16, 26). Vesicular stomatitis virus G glycoprotein (VSV-G)-pseudotyped Vif-proficient (WT) or Vif-deficient (ΔVif) viruses were generated in HEK293T cells by cotransfection with the corresponding full-length infectious molecular clone (IMC) and a VSV-G expression vector, and were used to infect parental and THP-1#11-4 cells at a multiplicity of infection (MOI) of 0.25 to establish virus-producing cells (**Fig. 2A**). Viral output was quantified by measuring HIV-1 p24 capsid levels in culture supernatants by enzyme-linked immunosorbent assay (ELISA) (**Fig. 2B, 2C**).

**Figure 2.**
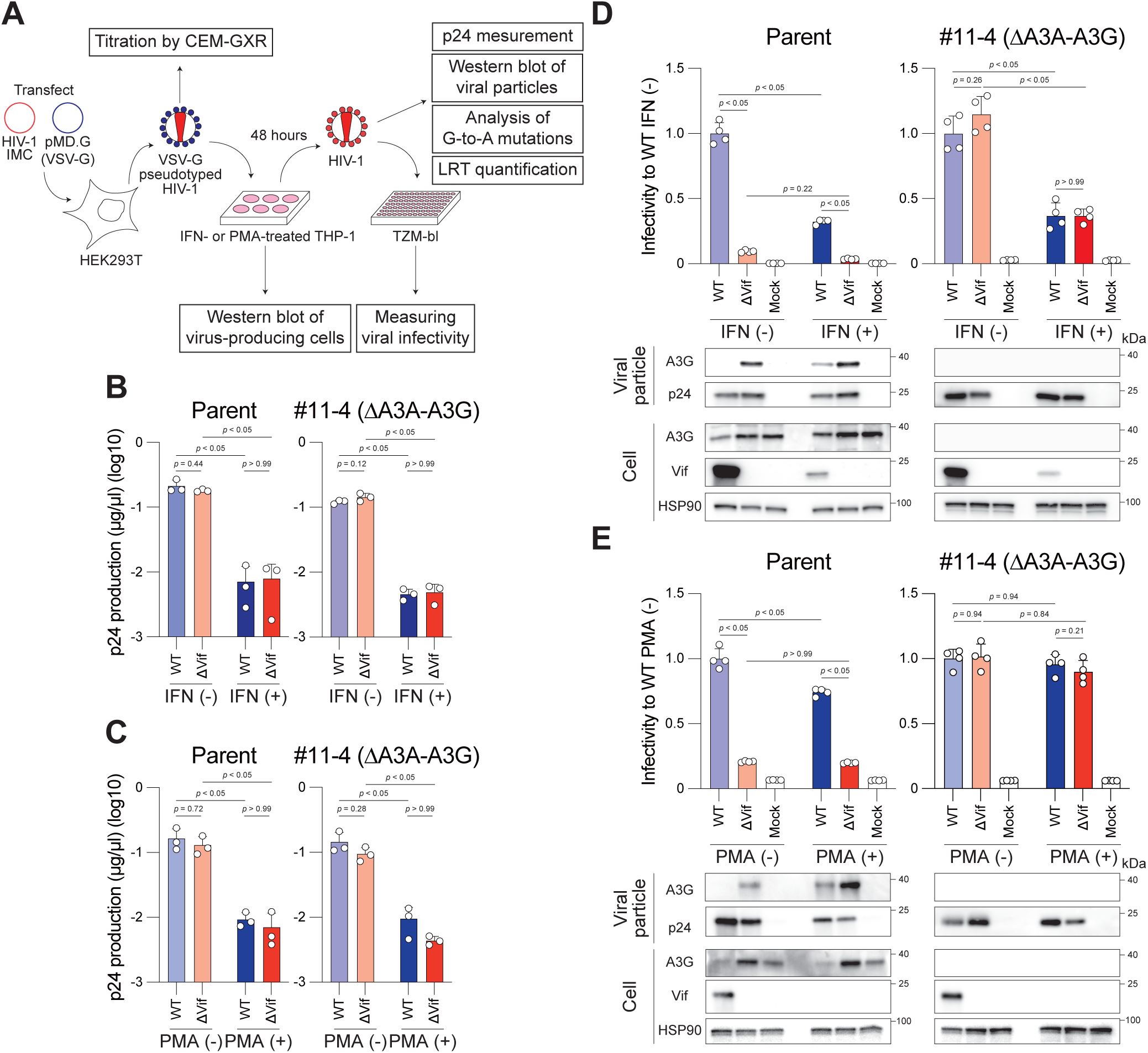
Effect of Vif on infectivity of HIV-1 produced from IFN- or PMA-treated parental and THP-1#11-4 cells. **(A)** Schematic of the pseudo-single cycle infectivity assay under IFN or PMA treatment. See Materials & Methods for details. **(B)** p24 production under type I IFN treatment. Cells were treated with 500 units/ml IFN for 6 hours, infected at an MOI of 0.25, and incubated for 48 hours. Viral supernatants were quantified by p24 ELISA. Bars represent mean ± SD of three independent experiments. Statistical significance was determined using two-way ANOVA with Tukey’s multiple comparisons test; p-values are indicated. **(C)** p24 antigen production under PMA treatment. Cells were treated with 20 ng/ml PMA for 20 hours prior to infection (MOI = 0.25). Analysis was performed as in (**B**). **(D)** Infectivity of IIIB WT and ΔVif viruses under IFN-treated conditions. Viral inputs from (**B**) were normalized by p24 and used to infect TZM-bl cells. Infectivity is expressed relative to WT virus produced under untreated conditions. Bars represent mean ± SD of four technical replicates from one representative experiment (n = 3 independent experiments). Statistical analysis was performed as in (**B**). Bottom panels show representative Western blot analyses of viral particles and whole-cell lysates probed with the indicated antibodies. p24 and HSP90 served as loading controls. **(E)** Infectivity of IIIB WT and ΔVif viruses under PMA-treated conditions. Viral inputs from (**C**) were normalized by p24 and used to infect TZM-bl cells. Data and statistical analysis are as described in (**D**).

Both IFN and PMA treatment reduced p24 production by more than 10-fold in both parental and THP-1#11-4 cells (**Fig. 2B**, **2C**). Importantly, WT and ΔVif viruses produced comparable levels of p24 under all conditions tested. These results indicate that IFN- and PMA-mediated suppression of viral output occurs independently of Vif and A3 family proteins.

### Vif is required for infectious virion production exclusively through antagonism of A3 proteins under IFN or PMA-stimulated conditions

We next assessed whether HIV-1 Vif is required to maintain virion infectivity under IFN- or PMA-stimulated conditions. Using pseudo-single cycle infectivity assays, infectious viruses were generated from untreated or stimulated parental and THP-1#11-4 cells, normalized by p24 levels, and analyzed for infectivity in TZM-bl cells, A3 incorporation, G-to-A mutation frequency, and late reverse transcription (LRT) products (**Fig. 2A**).

In parental THP-1 cells, ΔVif viruses exhibited markedly reduced infectivity compared to WT virus under unstimulated and stimulated conditions (**Fig. 2D**, **2E**). After p24 normalization, IFN reduced WT infectivity by approximately 50%, whereas PMA had minimal effect (**Fig. 2D**, **2E**). This difference is consistent with the possibility that IFN induces additional antiviral factor(s) that impair HIV-1 infectivity beyond the contribution of A3 family proteins [reviewed in (69–71)]. Nevertheless, ΔVif viruses remained profoundly defective under all conditions tested (**Fig. 2D**, **2E**).

In THP-1#11-4 cells, where *A3A*-*A3G* genes are genetically disrupted, infectivity of ΔVif virus was fully restored to WT levels under both unstimulated and stimulated conditions, including IFN treatment and PMA differentiation (**Fig. 2D**, **2E**). Although IFN reduced overall infectivity of both WT and ΔVif viruses, no difference was observed between genotypes (**Fig. 2D**, **2E**). These results demonstrate that the infectivity defect of ΔVif virus under stimulated conditions is entirely dependent on A3 family proteins.

Western blot analyses of A3G further supported this conclusion. In parental THP-1 cells, loss of Vif increased cellular A3G levels and its incorporation into virions under both untreated and stimulated conditions, correlating with reduced ΔVif infectivity (**Fig. 2D**, **2E**). In contrast, A3G was undetectable in THP-1#11-4 cells and in corresponding ΔVif virions, consistent with similar infectivity of WT and ΔVif viruses under both IFN- and PMA-treated conditions (**Fig. 2D**, **2E**).

Collectively, these data suggest that even under IFN-induced antiviral states or PMA-induced differentiation, the critical role of Vif in THP-1 cells is restricted to antagonism of A3 family proteins with no additional stimulation-induced Vif targets required for virion infectivity.

### A3 proteins are the exclusive source of stimulation-enhanced G-to-A mutations

The deaminase-dependent activity of A3 proteins, resulting in G-to-A mutations, represents the best-characterized mechanism underlying their antiviral function [reviewed in (1–4)]. To assess proviral mutagenesis under IFN or PMA-stimulated conditions, the *pol* region of HIV-1 proviral DNA recovered from SupT11 cells was amplified by nested PCR (**Fig. 2A**).

As expected, few G-to-A mutations were detected in proviruses of WT HIV-1 produced from untreated or IFN- or PMA-treated parental THP-1 cells (**Fig. 3A** to **3D**). In contrast, ΔVif viruses produced from parental THP-1 cells exhibited a marked increase in G-to-A mutations, predominantly in a GG-to-AG context (**Fig. 3A** to **3D**), consistent with antiviral activity of A3G protein (7, 8, 15, 16, 20–22, 24, 26). Notably, both IFN and PMA further increased the frequency of G-to-A mutations in ΔVif proviruses compared to untreated conditions (**Fig. 3A** to **3D**). Importantly, disruption of *A3A* through *A3G* abolished G-to-A mutations in ΔVif proviruses under both unstimulated and stimulated conditions (**Fig. 3A** to **3D**). These findings indicate that, even under IFN-induced antiviral states or PMA-induced differentiation, proviral G-to-A mutations are entirely dependent on A3 family proteins.

**Figure 3.**
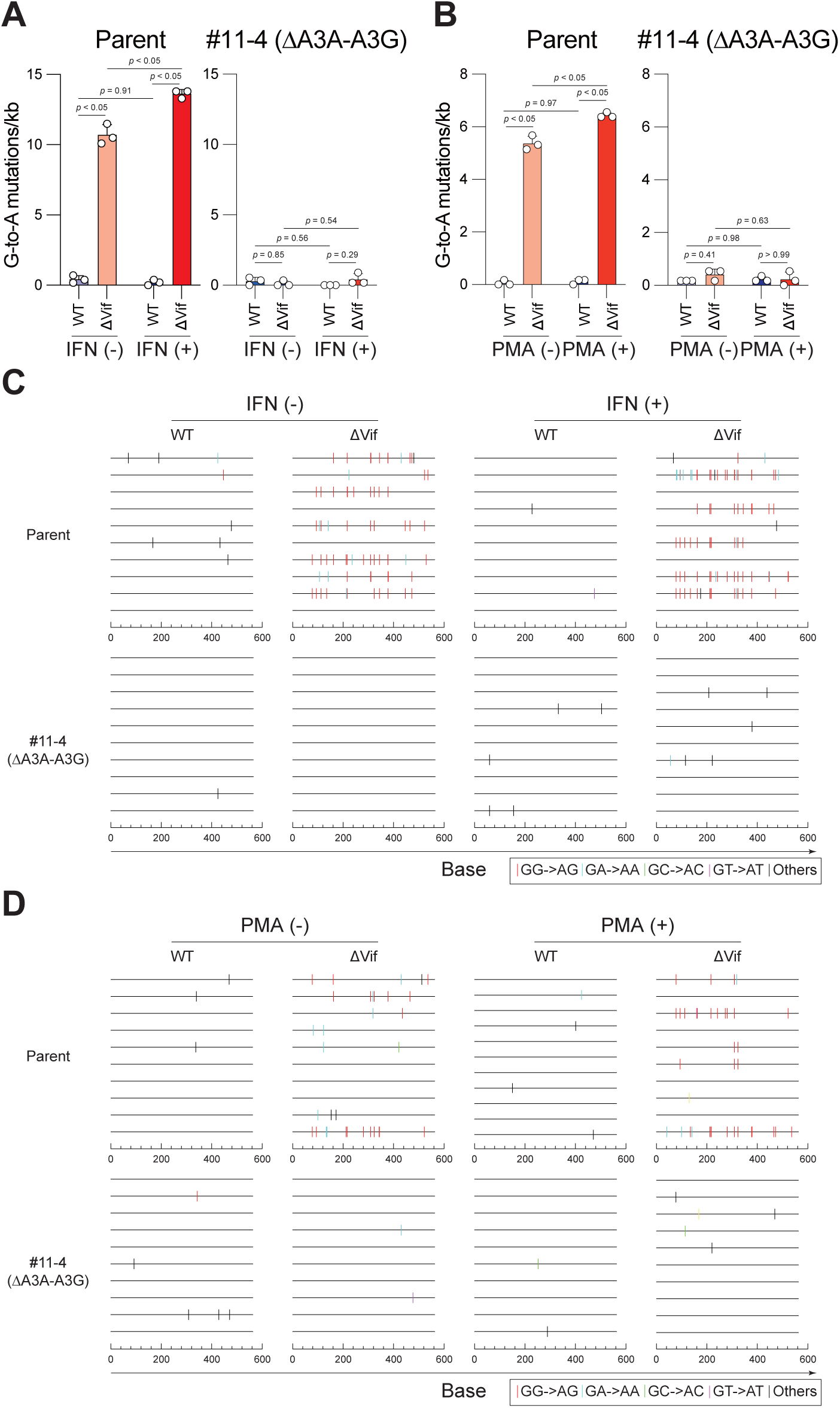
Effect of type I IFN or PMA treatment on deaminase-dependent A3 antiviral activity. (**A, B**) Quantification of G-to-A mutations in a 564-bp *pol* fragment following infection of SupT11 cells with WT or ΔVif viruses produced from parental or THP-1#11-4 cells under IFN **(A)** or PMA (**B**) conditions. Data represent mean ± SD of three independent experiments. Statistical analysis was performed using two-way ANOVA with Tukey’s test. (**C, D**) G-to-A mutation profiles showing dinucleotide contexts across the 564-bp *pol* amplicon for viruses produced under IFN (**C**) or PMA (**D**) conditions. Each vertical line indicates a mutation site categorized by dinucleotide context shown in the legend.

### IFN induces A3-independent reverse transcription blockade that is not antagonized by Vif

In addition to their deaminase-dependent mutagenic activity, A3 proteins can inhibit reverse transcription through deaminase-independent mechanisms [reviewed in (2, 3, 18, 19)]. We therefore asked whether HIV-1 Vif can counteract IFN-induced antiviral factors that restrict reverse transcription. To address this, HIV-1 LRT products were quantified by quantitative PCR (qPCR) (**Fig. 2A**).

In parental THP-1 cells, ΔVif viruses exhibited reduced LRT levels compared to WT virus under both unstimulated and IFN-stimulated conditions (**Fig. 4A**), consistent with A3-mediated restriction [reviewed in (1–3, 18, 19)]. Importantly, IFN treatment significantly reduced LRT product levels of WT HIV-1 by approximately 60% relative to untreated controls (**Fig. 4A**). A comparable reduction was observed for ΔVif virus under IFN treatment (**Fig. 4A**). Notably, IFN similarly reduced LRT levels of WT and ΔVif viruses even in THP-1#11-4 cells lacking A3A-A3G (**Fig. 4A**), indicating the presence of A3-independent reverse transcription blocks induced by IFN. Under these conditions, Vif expression did not restore LRT levels (**Fig. 4A**), demonstrating that Vif does not antagonize these IFN-induced restriction pathways. In contrast, PMA treatment had a minimal effect on LRT product levels of WT HIV-1 in either parental or THP-1#11-4 cells, and no significant differences were observed between WT and ΔVif viruses in THP-1#11-4 cells under either untreated or PMA-treated conditions (**Fig. 4B**).

**Figure 4.**
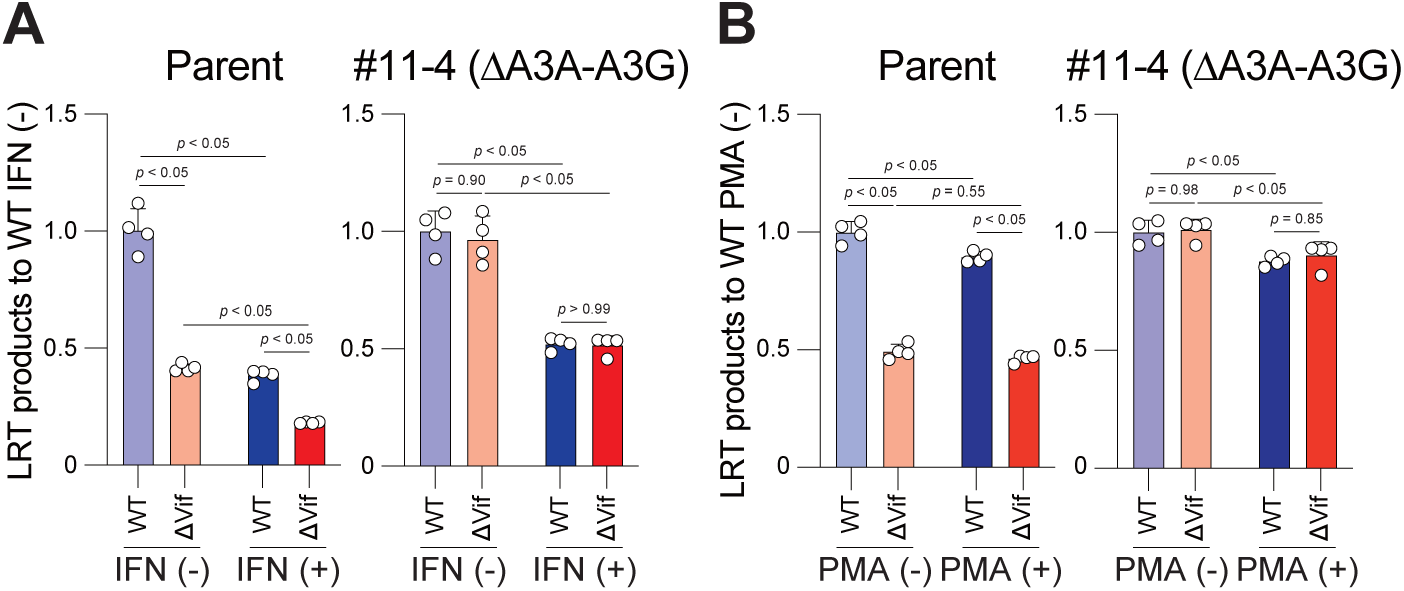
Effect of type I IFN or PMA treatment on deaminase-independent A3 antiviral activity. (**A, B**) Quantification of LRT products following infection of SupT11 cells with p24-normalized WT and ΔVif viruses (10 ng) produced from parental or THP-1#11-4 cells under IFN (**A**) or PMA (**B**) conditions. LRT products were quantified by qPCR, normalized to *CCR5* gene copy number, and expressed relative to WT virus under untreated conditions. Bars represent mean ± SD of four technical replicates. The data shown are representative of three independent experiments. Statistical significance for each condition was determined using two-way ANOVA with Tukey’s multiple comparisons test; p-values are indicated on the graphs.

Together, these findings demonstrate that IFN induces A3-independent restriction at the level of reverse transcription that is not counteracted by Vif, highlighting the specificity of Vif for A3-mediated antiviral pathways.

### A3-dependent Vif function is conserved across diverse HIV-1 strains under IFN- or PMA-stimulated conditions

To determine whether A3-dependent Vif function is conserved across HIV-1 strains, we examined multiple HIV-1 strains, including transmitted/founder (TF) HIV-1 subtype B and C viruses. WT and ΔVif viruses were produced in either parental or THP-1#11-4 cells under IFN or PMA-treated conditions, and viral infectivity was assessed using TZM-bl cells.

Consistent with the results obtained using the IIIB strain (**Fig. 2D**, **2E**), all ΔVif viruses exhibited markedly reduced infectivity in parental THP-1 cells compared to their corresponding WT viruses under both IFN and PMA stimulation (**Fig. 5**, **Fig. S1**). In contrast, disruption of *A3A*-*A3G* in THP-1#11-4 cells fully restored ΔVif infectivity to WT levels across all strains tested (**Fig. 5**, **Fig. S1**). Notably, the LAI strain exhibited complete rescue of ΔVif infectivity despite its Vif being defective in PPP2R5A degradation (47). Together, these findings demonstrate that Vif-dependent infectivity in THP-1 cells is universally mediated through antagonism of A3 proteins across diverse HIV-1 strains and stimulation conditions.

**Figure 5.**
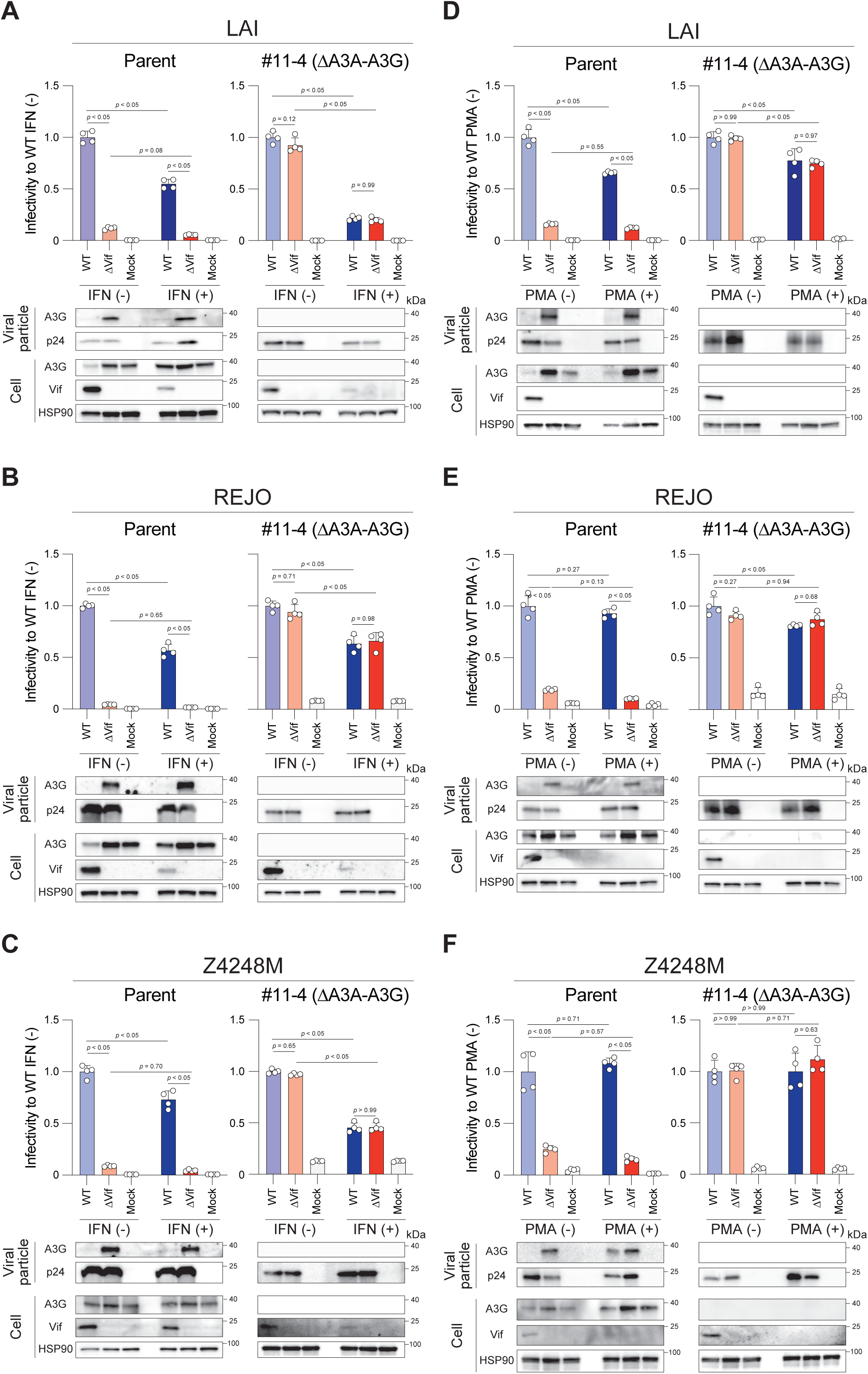
Vif-dependent infectivity of diverse HIV-1 strains under IFN- or PMA-treated conditions. Infectivity of laboratory-adapted [LAI (**A**, **D**)], TF subtype B [REJO (**B**, **E**)], or TF subtype C [Z4248M (**C**, **F**)] viruses produced under IFN (**A** to **C**) or PMA (**D** to **F**) treatment. Viral inputs were normalized by p24 and used to infect TZM-bl cells. Infectivity was expressed relative to WT virus produced under untreated conditions. Bars represent mean ± SD of four technical replicates from one representative experiment (n = 3 independent experiments). Statistical significance was determined using two-way ANOVA with Tukey’s test. Bottom panels show representative Western blot analyses of viral particles and whole-cell lysates probed with the indicated antibodies. p24 and HSP90 served as loading controls.

## Discussion

HIV-1 Vif is best known for antagonizing the antiviral activity of A3 proteins; however, it has also been reported to degrade additional host factors, including components of cell cycle regulatory pathways (45–50). It has remained unclear whether such interactions make an essential contribution to infectious virion production, particularly under inflammatory or differentiation conditions. In this study, we directly addressed this question using IFN-stimulated and PMA-differentiated THP-1 monocytic cells. Our data demonstrate that, even under these conditions, the essential role of Vif in maintaining virion infectivity is fully explained by antagonism of A3 proteins.

Type I IFN induces a broad antiviral state, upregulating numerous ISGs, including several A3 family members and antiviral factors such as Tetherin and IFITMs that restrict HIV-1 at multiple stages of replication [reviewed in (69–71)]. Consistent with this, we observed substantial suppression of viral production independent of Vif (**Fig. 2B**). Notably, even after p24 normalization, IFN reduced WT virion infectivity, whereas PMA had little effect (**Fig. 2D, 2E**), suggesting that IFN induces additional antiviral factor(s) that impair HIV-1 infectivity beyond the contribution of A3 family proteins. Importantly, however, these IFN-induced effects did not uncover any additional essential Vif-dependent pathway required for maintenance of infectivity. ΔVif viruses produced in IFN-treated parental cells exhibited severe infectivity defects, yet complete disruption of *A3A*-*A3G* fully restored infectivity under all conditions tested (**Fig. 2D**). Moreover, although IFN induced blocks at the level of reverse transcription, Vif did not antagonize these A3-independent restrictions (**Fig. 4A**), highlighting the functional specificity of the Vif-A3 axis.

Vif-mediated degradation of PPP2R5 family proteins and associated cell cycle perturbations have been proposed to represent additional functional outputs of Vif (45–50). Importantly, the LAI strain exhibited complete rescue of ΔVif infectivity despite its inability to degrade PPP2R5A (47) (**Fig. 5A**, **5D**), indicating that this interaction is dispensable for infectious virion production in this single cycle infection model. Although these pathways may influence cellular physiology or multi-cycle replication dynamics, they do not constitute essential determinants of virion infectivity under the conditions examined here.

Notably, no factor other than A3 proteins contributed to detectable G-to-A mutations under any condition tested, including untreated, IFN-stimulated, and PMA-differentiated conditions (**Fig. 3A, 3C**). These results reinforce the unique role of A3-mediated mutagenesis as the irreversible genomic threat that Vif must neutralize to preserve infectivity. In contrast, IFN-induced antiviral factors restrict HIV-1 through non-mutagenic mechanisms and therefore do not impose the same irreversible genetic damage on the viral genome.

The inability of inflammatory stimulation or differentiation to reveal additional essential Vif targets suggests that evolutionary pressure on Vif is predominantly driven by the need to evade A3-mediated mutagenesis. A3 enzymes impose lethal or highly deleterious mutations on viral genomes [reviewed in (1–3)], creating an irreversible genetic threat that directly compromises infectivity. Thus, while IFN-induced, A3-independent antiviral pathways can limit viral replication, they do not create the same selective pressure for Vif-mediated counteraction. Consistent with this, strict A3 dependence of Vif was conserved across laboratory-adapted and TF HIV-1 strains (**Fig. 5**, **Fig. S1**), indicating that this functional architecture is maintained across diverse viral genetic backgrounds. Together, these findings support the conclusion that antagonism of A3 proteins constitutes the core and indispensable function of Vif.

Several limitations should be noted. Our analyses were performed in a monocytic cell line and primarily in single-cycle infection assays. It remains possible that primary macrophages, multi-cycle replication systems, or *in vivo* inflammatory environments may reveal context-dependent Vif activities not captured here. Nevertheless, within the defined experimental framework, A3 antagonism fully accounts for the essential role of Vif in maintaining infectious virion production.

In summary, antagonism of A3 family proteins fully accounts for the essential role of Vif in infectious HIV-1 production in THP-1 monocytic cells, irrespective of IFN stimulation or differentiation state. These findings indicate that the core function of Vif is neutralization of A3-mediated restriction and mutagenesis. Although further studies in primary CD4⁺ T lymphocytes and macrophages will be required to define the generalizability, our data suggest that evolutionary pressure on Vif is driven predominantly by evasion of A3-imposed genomic damage. Targeting Vif may therefore expose HIV-1 to irreversible intrinsic mutagenesis even under inflammatory conditions.

## Materials & Methods

### Cell lines and culture

Human embryonic kidney 293T (HEK293T) [American Type Culture Collection (ATCC), Cat# CRL-3216], TZM-bl (72) [NIH AIDS Reagent Program (NARP), Cat# ARP-8129], and HeLa-CD4 (73) (NARP, Cat# ARP-154) were maintained in high-glucose Dulbecco’s Modified Eagle Medium (DMEM) (Wako, Cat# 044-29765) supplemented with 10% fetal bovine serum (FBS) (NICHIREI, Cat#175012) and 1% Penicillin/Streptomycin (P/S) (Wako, Cat# 168-23191). Parental THP-1 cells (53) were provided by Dr. Andrea Cimarelli (INSERM, France), CEM-GXR cells (74) were provided by Dr. Todd Allen (Harvard University, USA), and the *A3*-null SupT1 derivative SupT11 cells (75) were provided by Dr. Reuben Harris (University of Texas Health San Antonio, USA). The establishment of THP-1#11-4 cells has been described previously (16). These suspension cell lines were cultured in Roswell Park Memorial Institute (RPMI) 1640 medium (Thermo Fisher Scientific, Cat# C11875500BT) supplemented with 10% FBS and 1% P/S.

### Plasmids

Full-length Vif-proficient and Vif-deficient (X^26^ and X^27^) HIV-1 IIIB C200 and LAI proviral expression constructs were described previously (31, 76). The full-length Vif-proficient NL-CSFV3 proviral expression plasmid was kindly provided by Dr. Kei Sato (University of Tokyo, Japan) (77). The following full-length TF HIV-1 infectious molecular clones were obtained through the NARP: REJO (Cat# ARP-11746), RHPA (Cat# HRP-11744), Z3618M (Cat# ARP-13268), and Z4248M (Cat# ARP-13277).

Vif-deficient HIV-1 mutants were generated by introducing double stop codons at amino acid positions 26 and 27 in the *vif* gene using primers listed in **Table S1** (TI collection numbers correspond to the oligonucleotides used in this study). Site-directed mutagenesis was performed for the following clones: NL-CSFV3 (TI351), REJO (TI497), RHPA (TI487), Z3618M (TI467), and Z4248M (TI536 and TI537). DNA fragments containing the introduced *vif* mutations were amplified using the primer sets indicated in **Table S1** (NL-CSFV3: TI347/TI348 and TI349/TI350, REJO: TI493/TI494 and TI495/TI496, RHPA: TI483/TI484 and TI485/TI486, Z3618M: TI463/TI464 and TI465/TI466, and Z4248M: TI535 and TI538) and inserted into the corresponding HIV-1 infectious molecular clone using the In-Fusion® Snap Assembly Master Mix (Takara Bio, Cat# 638948). All inserted sequences were verified by Sanger sequencing (AZENTA) using the primers listed in **Table S1** (NL-CSFV3: TI362/TI363/TI364/TI365, REJO: TI519/TI520/TI521/TI522/TI523, RHPA: TI488/T489/TI490/TI491/TI492, Z3618M: TI481/TI482, and Z4248M: TI539/TI540), and sequence data were analyzed with Sequencher v5.4.6 (Gene Codes Corporation).

### RT-qPCR

Cells were harvested by centrifugation and washed with phosphate-buffered saline (PBS). Total RNA was extracted using an RNA isolation kit (NIPPON Genetics, Cat# FG-81250). Complementary DNA (cDNA) was synthesized from the purified RNA using Transcriptor Reverse Transcriptase (Roche, Cat# 03531287001) with random primers. RT-qPCR was then performed using Power SYBR Green PCR Master Mix (Thermo Fisher Scientific, Cat# 4367659). Primer sequences for each *A3* mRNA were designated based on a previous report (67). Primer sets for ISGs used in RT-qPCR are listed in **Table S1**. Fluorescence signals were detected using a Thermal Cycler Dice Real Time System III (Takara Bio), and mRNA levels were normalized to *TATA-binding protein* (*TBP*) mRNA levels.

### Pseudo-single cycle infectivity assay

Single-round HIV-1 infection experiments using VSV-G-pseudotyped viruses were performed as previously described (15, 16, 26) (**Fig. 2A**). To produce VSV-G-pseudotyped infectious viruses, HEK293T cells (2 × 10^6^) were transfected with 2.4 µg of an HIV-1 IMC and 0.6 µg of a VSV-G expression plasmid (Addgene #138479) using TransIT-LT1 (Takara, Cat# MIR2306). After 48 hours, culture supernatants containing VSV-G-pseudotyped viruses were harvested and filtered through 0.45 µm membranes (Merck, Cat# SLHVR33RB). To determine the viral infectivity, the filtered supernatant was used to infect CEM-GXR cells (2.5 × 10^4^). After 48 hours, GFP-positive cells were quantified by flow cytometry (CytoFLEX, Beckman Coulter) and analyzed with FlowJo software v10.7.1 (BD Biosciences).

For virus production, THP-1 cells were pretreated with either 500 units/ml of type I IFN (PBL Assay Science, Cat# 11200-2) for 6 hours or 20 ng/ml of PMA (Selleck, Cat# S7791) for 20 hours, followed by washing with PBS. Cells were then infected with VSV-G-pseudotyped viruses at an MOI of 0.25. At 24 hours post-infection, cells were washed twice with PBS to remove residual virus and cultured for an additional 24 hours. Supernatants were collected, filtered through 0.45 µm membranes, and analyzed for p24 antigen concentration using the p24 ELISA kit 2.0 (ZeptoMetrix, Cat# 0801008).

Equal amounts of p24-normalized viruses (0.5–5 ng of p24 antigen) were subsequently used to infect TZM-bl cells (1 × 10^4^) seeded in 96-well white plates (Sumitomo Bakelite, Cat# MS-8096W). After 48 hours, the culture medium was removed and luciferase activity was measured using the Bright-Glo Luciferase Assay System (Promega, Cat# E2620) with a Centro XS3 LB963 microplate luminometer (Berthold Technologies).

### Western blotting

Western blotting was performed as previously described (15, 16, 78). Briefly, virus-producing cells were harvested, washed with PBS, and lysed in lysis buffer [25 mM HEPES (pH 7.2), 10% Glycerol, 125 mM NaCl, 1% NP40] containing Nonidet P40 (NP40) (Nacalai Tesque, Cat# 18558-54). Cell lysates were incubated on ice for 30 minutes and clarified by centrifugation at 22,000 × *g* for 15 minutes at 4°C. Protein concentrations were determined using the Bio-Rad Protein Assay Dye Reagent Concentrate (Bio-Rad, Cat# 5000006). Lysates were mixed with 2 × sodium dodecyl sulfate (SDS) sample buffer [100 mM Tris-HCl (pH 6.8), 4% SDS, 20% glycerol, 0.05% bromophenol blue] containing 12% β-mercaptoethanol (2-ME) and boiled at 95°C for 5 minutes.

For analysis of virion-associated proteins, viral particles were pelleted by underlaying 1 ml of viral supernatant onto 500 µl of 20% sucrose/PBS and centrifuging at 22,000 × *g* for 2 hours at 4°C. The resulting pellets were resuspended in 50 µl of 2 × SDS sample buffer containing 12% 2-ME and boiled at 95°C for 5 minutes.

Cell lysates (1-5 µg of total protein) or virus lysates (1–10 ng of p24 antigen) were resolved by 12% SDS–polyacrylamide gel electrophoresis (SDS-PAGE) and transferred to PVDF membranes (Millipore, Cat# IPVH00010). Membranes were blocked for 1 hour at room temperature in Tris-buffered saline (TBS) (25 mM Tris, 137 mM NaCl, 2.68 mM KCl) solution containing 4% skim milk (Wako, Cat# 190-12865) and 0.1% Tween-20 (Bio-Rad, Cat# 1610781) (0.1% TBST).

Primary antibodies were diluted in 4% skim milk/0.1% TBST and incubated with membranes overnight at 4°C. The primary antibodies and their dilutions were as follows: mouse anti-HSP90 (BD Transduction Laboratories, Cat# 610418, 1:5,000); rabbit anti-A3G (NARP, Cat# ARP-10201, 1:2,500); mouse anti-Vif (NARP, Cat# ARP-6459, 1:2,000 used for IIIB, LAI, REJO, and NL-CSFV3); rabbit anti-Vif (NARP, Cat# ARP-809, 1:2,000 used for RHPA and Z4248M); and mouse anti-p24 (NARP, Cat# ARP-1513, 1:2,000).

After three washes with 0.1% TBST (10 minutes each), membranes were incubated for 1 hour at room temperature with horseradish peroxidase (HRP)-conjugated secondary antibodies diluted in 1% skim milk/0.1% TBST as follows: donkey anti-rabbit IgG polyclonal antibody (Jackson ImmunoResearch, Cat# 711-035-152; 1:5,000); donkey anti-mouse IgG polyclonal antibody (Jackson ImmunoResearch, Cat# 715-035-150; 1:5,000).

Following three additional washes in 0.1% TBST (10 minutes each), signals were developed using SuperSignal™ West Femto Maximum Sensitivity Substrate (Thermo Fisher Scientific, Cat# 34095) or SuperSignal™ West Atto Ultimate Sensitivity Substrate (Thermo Fisher Scientific, Cat# A38555), and visualized with an Amasham™ Imager 600 (Cytica) or qTouch Western Blot imager (RWD Life Science).

### Hypermutation analysis

The method for analyzing mutations induced by A3 proteins was described previously (13, 15, 16, 26). Viruses obtained from the Pseudo-single cycle infectivity assay were used to infect SupT11 cells (5 × 10^5^), which were subsequently cultured for 48 hours. At 48 hours post-infection, cells were harvested and washed twice with PBS. Total DNA was extracted using the DNeasy Blood & Tissue Kit and treated with RNase A according to the manufacturer’s instructions. The purified DNA was digested with DpnI, and the *pol* region of HIV-1 was amplified by nested PCR using outer primers (TI150/TI151; 876 bp product) and inner primers (TI152/TI153; 564 bp product). PCR products were purified with a Gel Extraction Kit (Thermo Fisher Scientific, Cat# K0692) and cloned into the pJET1.2/blunt vector (Thermo Fisher Scientific, Cat# K1232). Plasmid inserts were sequenced using the pJET forward primer (listed in **Table S1**) by Sanger sequencing (AZENTA) and analyzed using Sequencher v5.4.6. At least ten independent clones were analyzed for each condition. Hypermutation analysis was performed using the HIV Sequence Database Hypermut tool (https://www.hiv.lanl.gov/content/sequence/HYPERMUT/hypermut.html). Clones exhibiting identical mutation patterns were excluded from the analysis.

### Quantification of LRT products

To quantify the level of LRT products, viral particles were first produced by infecting THP-1 cells with VSV-G-pseudotyped HIV-1 (see details in the **Pseudo-single cycle infectivity assay** section). The resulting viral particles were quantified by p24 ELISA. Next, HeLa-CD4 cells (2 × 10^5^) were infected with viral supernatants containing 10 ng of p24 antigen. At 12 hours post-infection, cells were harvested and washed twice with PBS. Total DNA was extracted using the DNeasy Blood & Tissue Kit (Qiagen, Cat# 69504) and treated with RNase A (Qiagen, Cat# 19101) according to the manufacturer’s instructions. After digestion with DpnI (New England BioLabs, Cat# R0176S), 50 ng of purified DNA was used for qPCR to amplify both the LRT products and the *CCR5* reference gene. Primer sequences used for qPCR were as follows: TI122 andTI123 for LRT, and TI124 and TI125 for *CCR5* (**Table S1**). qPCR was performed using Power SYBR Green PCR Master Mix (Thermo Fisher Scientific, Cat# 4367659), and fluorescence signals were acquired with a Thermal Cycler Dice Real Time System III (Takara Bio).

### Statistical analysis

Statistical analyses were performed using an unpaired t-test with Welch’s correction (**Fig. 1**) or two-way ANOVA followed by Tukey’s multiple comparisons test (**Fig. 2B** to **2E, 3A, 3B, 4A, 4B, 5, Fig. S1**). All statistical tests were conducted using GraphPad Prism software v8.4.3.

## Acknowledgements

We would like to thank all members of the Ikeda lab for their supports. We also thank Drs. Todd Allen, Andrea Cimarelli, Reuben Harris, and Kei Sato for generously sharing reagents. This study was supported in part by the AMED Research Program on HIV/AIDS (JP22fk0410055, to TI; JP25fk0410065h0002, to KM); the AMED Research Program, HK2-MIRAI (JP256f0137011, to TI); the JSPS KAKENHI Grant-in-Aid for Scientific Research C (22K07103, to TI); the JSPS Leading Initiative for Excellent Young Researchers (LEADER) (to TI); the Mitsubishi Foundation (to TI), the Takeda Science Foundation (to TI); the Uehara Memorial Foundation (to TI); the Heiwa Nakajima Foundation (to TI); the G-7 Scholarship Foundation (to TI); the International Joint Research Project of the Institute of Medical Science, the University of Tokyo (to TI); the Joint Usage/Research Center on Tropical Disease, Institute of Tropical Medicine, Nagasaki University (2026-Ippan-12, to TI); and the scholarship program of The Chemo-Sero-Therapeutic Research Institute (KAKETSUKEN) (to RS).

The authors declare that they have no competing interests.

---

**Supplemental Figure 1. Vif-dependent infectivity of additional HIV-1 strains under IFN-or PMA-treated conditions.**

Infectivity of laboratory-adapted [NL-CSFV3 (**A**, **D**)], TF subtype B [RHPA (**B**, **E**)], or TF subtype C [Z3618M (**C**, **F**)] viruses produced under IFN (**A** to **C**) or PMA (**D** to **F**) treatment. Viral inputs were normalized by p24 and used to infect TZM-bl cells. Infectivity was expressed relative to WT virus produced under untreated conditions. Bars represent mean ± SD of four technical replicates from one representative experiment (n = 3 independent experiments). Statistical significance was determined using two-way ANOVA with Tukey’s test. Bottom panels show representative Western blot analyses of viral particles and whole-cell lysates probed with the indicated antibodies. Notably, Z3618M Vif was not detected using either mouse monoclonal or rabbit polyclonal anti-Vif antibodies employed in this study. p24 and HSP90 served as loading controls.

